# Node Detection using High-Dimensional Fuzzy Parcellation Applied to the Insular Cortex

**DOI:** 10.1101/011999

**Authors:** Ugo Vercelli, Matteo Diano, Tommaso Costa, Sergio Duca, Giuliano Geminiani, Alessandro Vercelli, Franco Cauda

**Author notes:** Corresponding author: Dr. Tommaso Costa, Department of Psychology, Via Po 14, 10123 Turin, Italy.

## Abstract

Several functional connectivity approaches require the definition of a set of ROIs that act as network nodes. Different methods have been developed to define these nodes and to derive their functional and effective connections, most of which are rather complex. Here we aim to propose a relatively simple “one-step” border detection and ROI estimation procedure employing the fuzzy c-mean clustering algorithm.

To test this procedure and to explore insular connectivity beyond the two/three-region model currently proposed in the literature, we parcellated the insular cortex of a group of twenty healthy right-handed volunteers (10 females) scanned in a resting state condition.

Employing a high-dimensional functional connectivity-based clustering process, we confirmed the two patterns of connectivity previously described. This method revealed a complex pattern of functional connectivity where the two previously detected insular clusters are subdivided into several other networks, some of which not commonly associated with the insular cortex, such as the default mode network and parts of the dorsal attentional network. Finally, the detection of nodes was reliable as demonstrated by the confirmative analysis performed on a replication group of subjects.

## INTRODUCTION

One powerful method for studying brain organization is the graph-theoretic approach (Bullmore and Sporns 2009). Like with other connectivity methods such as seed-based functional connectivity and diffusion tensor imaging approaches, this method requires the definition of a set of ROIs that act as network nodes (Smith et al. 2011). Several techniques have been employed to functionally derive these nodes and their connections (for example see (Nelson et al. 2010a; Cohen et al. 2008) The definition of such nodes often involves complicated functional connectivity estimation and border detection procedures. Here, we aim to propose a relatively simple “one-step” border detection and ROI estimation procedure. Indeed, we propose to take advantage of one of the characteristics of the fuzzy c-mean clustering algorithm (Cauda et al. 2010): this procedure allows a fixed percent of voxels with a borderline pattern of connectivity to be non-unequivocally attributed. The further we move from the center towards the border of a cluster, the more the characteristics of the pattern of connectivity are intermixed with those of neighboring clusters (e.g. non-unequivocally determined); indeed as recently evidenced by Smith et al. (Smith et al. 2012) the maximization of the spatial independence could lead to a suboptimal detection of networks that shares significant spatial overlaps. Our methods maximizes the temporal independence at the expense of a percent of voxels non univocally attributed.

Previous clustering studies such as those by Cauda et al. (Cauda et al. 2011; Cauda et al. 2010; Chang et al. 2012; Deen et al. 2011) performed the clustering procedure at subject level. Given the relative deficiency of time points (about 120-200, in a 6-minute run), this procedure only has good reliability for low-dimensional parcellation (e.g. with a limited number of clusters). In this study, as suggested by others (Kiviniemi et al. 2009; Leech et al. 2012; Smith et al. 2011) we concatenated the time courses across all subjects to constitute a very big dataset that allowed us to obtain higher clustering dimensionality. Rather than being a step backward, this “fixed-effect” approach permits a good estimation of a common set of clusters (and thus nodes) for a given group of subjects; between-subject variance was taken into consideration in the subsequent step, where each node’s functional connectivity pattern was evaluated at the subject level and then summarized using a random-effect analysis. This method is actually very similar to the dual regression approach (Zuo et al. 2010) where Independent Component Analysis was first applied to the concatenated dataset. To investigate how this method performs with real data, we applied our procedure to the insular surface of a group of twenty healthy subjects scanned in a resting state condition. A second dataset involving eighteen volunteers was used for replication testing. We chose the insular surface because the insula is a complex and pivotal (Nelson et al. 2010b) brain area where different inputs from the body and the external world are integrated (Craig 2009); this area has been parcellated using different measures such as resting state functional connectivity (Cauda et al. 2011; Deen et al. 2011; Nelson et al. 2010b), task-related functional connectivity (Cauda et al. 2012a; Kurth et al. 2010b) and Diffusion Tensor Imaging (Nanetti et al. 2009; Cerliani et al. 2011) in two (Cauda et al. 2012a; Cauda et al. 2011; Cloutman et al. 2012; Jakab et al. 2012; Taylor et al. 2009), three(Chang et al. 2012; Deen et al. 2011) or more clusters (Cohen et al. 2008; Kurth et al. 2010b; Power et al. 2011; Yeo et al. 2011) each of which with a unique pattern of connectivity. A recent paper by Kelly et al. (Kelly et al. 2012) demonstrated a convergence between resting state, task-based functional connectivity and anatomical coactivations at several different parcellation levels (from 2 to 12 clusters per side, with more than 12 clusters reliability dropped by about 50%) thus supporting a common hierarchical structure of the insular cortex, demonstrated through different modalities.

Given the relevance of this brain area and to confirm these recent findings, it would be of great interest to apply this new method to obtain a high-dimensional parcellation of the insular surface. Indeed, we suspect that high-dimensional clustering (Kiviniemi et al. 2009) will enable us to demonstrate the existence of a more complex pattern than in our previous studies (Cauda et al. 2012a; Cauda et al. 2011), with “echoes” (Leech et al. 2012) of several brain networks nested within the two main insular patterns previously reported.

## MATERIALS AND METHODS

### SUBJECTS

#### Main group

Twenty healthy right-handed volunteers (10 females, mean age 32.6 ± 11.2).

#### Replication group

Eighteen healthy right-handed volunteers (9 females, mean age 25.3 ± 4.2). All subjects were free of neurological or psychiatric disorders, not taking any medication known to alter brain activity, and had no history of drug or alcohol abuse. Handedness was ascertained with the Edinburgh Inventory (Oldfield 1971). We obtained the written informed consent of each subject, in accordance with the Declaration of Helsinki. The study was approved by the institutional committee on ethical use of human subjects at the University of Turin.

### TASK AND IMAGE ACQUISITION

Images were acquired during a resting state scan on a 1.5 Tesla INTERA™ scanner (Philips Medical Systems). Functional T2^*^ weighted images were acquired using echoplanar (EPI) sequences, with a repetition time (TR) of 2000 ms, an echo time (TE) of 50 ms, and a 90° flip angle. The acquisition matrix was 64 x 64, with a 200 mm field of view (FoV). A total of 200 volumes were acquired, with each volume consisting of 19 axial slices; slice thickness was 4.5 mm with a 0.5 mm gap; in-plane resolution was 3.1 mm. Two scans were added at the beginning of functional scanning to reach steady-state magnetization before acquiring the experimental data. A set of three-dimensional high-resolution T_1_-weighted structural images was acquired, using a Fast Field Echo (FFE) sequence, with a 25 ms TR, an ultra-short TE, and a 30° flip angle. The acquisition matrix was 256 x 256, and the FoV was 256 mm. The set consisted of 160 contiguous sagittal images covering the whole brain.

### DATA ANALYSIS

The datasets were pre-processed and analyzed using the BrainVoyager QX software (Brain Innovation, Maastricht, The Netherlands).

Functional images were pre-processed as follows to reduce artifacts (Miezin et al. 2000): (i) slice scan time correction was performed using a sinc interpolation algorithm; (ii) 3D motion correction was applied: using a trilinear interpolation algorithm, all volumes were spatially aligned to the first volume by rigid body transformations and the roto-translation information was saved for subsequent elaborations; (iii) spatial smoothing was performed using a Gaussian kernel of 8 mm FWHM.

Several nuisance covariates were regressed out from the time courses to control for the effects of physiological processes (such as fluctuations related to cardiac and respiratory cycles) (Bandettini and Bullmore 2008) (Birn et al. 2008) (Napadow et al. 2008) and motion. Specifically, we included 9 additional covariates from white matter (WM), Global signal (GS) (Fox et al. 2009) and cerebrospinal fluid (CSF), as well as 6 motion parameters.

The time courses were then i) temporally filtered in order to only keep frequencies between 0.008 and 0.08 Hz, ii) normalized.

After pre-processing, the datasets of each subject were transformed into Talairach space (Talairach and Tournoux 1988): the cerebrum was translated and rotated into the anterior-posterior commissure plane and its borders identified. Finally, using the anatomo-functional coregistration matrix and the determined Talairach reference points, the functional time course of each subject was transformed into Talairach space and the volume time course created.

The clustering procedure was the one described in Cauda et al. (Cauda et al. 2012b). In brief, we applied a fuzzy c-mean algorithm to the time courses of all the insular voxels and clustered these voxels on the basis of the temporal similarity between them. Unlike in the study by Cauda (Cauda et al. 2011): i) time courses were concatenated across all subjects; this step, as pointed out by others (Smith et al. 2011), makes it possible to obtain a much higher clustering dimensionality (i.e. more clusters); ii) in the visualization step we only visualized the 20% of voxels that, due to the fuzziness coefficient employed, were classified as non-unequivocal. Indeed, we considered these as border voxels, where the time course of the voxels shows transitional characteristics between contiguous clusters. The final step of this procedure was to place a spherical ROI with a radius of 3 mm in the local maxima of each cluster (i.e. the area of maximal similarity between voxel time courses). The optimal number of clusters (Bowman et al. 2004) was chosen using the Silhouette validation method (Rousseeuw 1987).

We employed a variant of the dual regression approach (Zuo et al. 2010) used also in (Leech et al. 2011) to investigate the specific pattern of connectivity of each cluster. In brief, we included all the subject-specific time courses of all the 12 right insular spherical ROIs in a multiple regression analysis. This method resulted in a subject-specific time course for each ROI, controlling for the variance explained by all the other ROIs (Leech et al. 2012).

To confirm previous connectivity results (Cauda et al. 2011) we also calculated the functional connectivity of the anterior and posterior insular clusters by grouping together the ROIs belonging to each cluster described in (see Fig 1 upper panel).

All maps were thresholded at p<0.05 and corrected for multiple comparisons using the False discovery rate (FDR).

## RESULTS

Our method separated 12 clusters for each insula: this number resulted as the preferred number of clusters from the application of the silhouette validation method (Rousseeuw 1987). The algorithm returned the borders of the functionally homogeneous areas; for each cluster a spherical ROI with a radius of 3 mm was placed in the area with the maximal homogeneity (see Fig 1 upper panel). The results of our method were replicable; indeed, as shown in the lower panel of Fig 1, all the ROIs were also found in the control group and the locations were almost overlapping in 17 out of 24 ROIs, while the other ROIs were displaced by only a few millimeters.

Our calculation of the functional connectivity of the ROIs belonging to anterior and posterior insular clusters confirmed the two patterns of connectivity previously described (Cauda et al. 2011). These two patterns broken down into low-dimensional ones prevail over all the others. The anterior pattern that occupies the anterior-ventralmost part of the insular cortex is characterized by a cingulate-frontoparietal pattern of connectivity which has often been related to salience detection; the posterior pattern that occupies the dorsal posterior insula shows a sensorimotor connectivity pattern. However, by considering all the 12 ROIs and including the ROI time courses in a multiple regression analysis (Leech et al. 2012) we discovered a much more intricate image. Using this method we demonstrated that these areas are connected to several other networks such as the default mode network, sensorimotor network and parts of the dorsal attentional network.

**Fig. 1.**
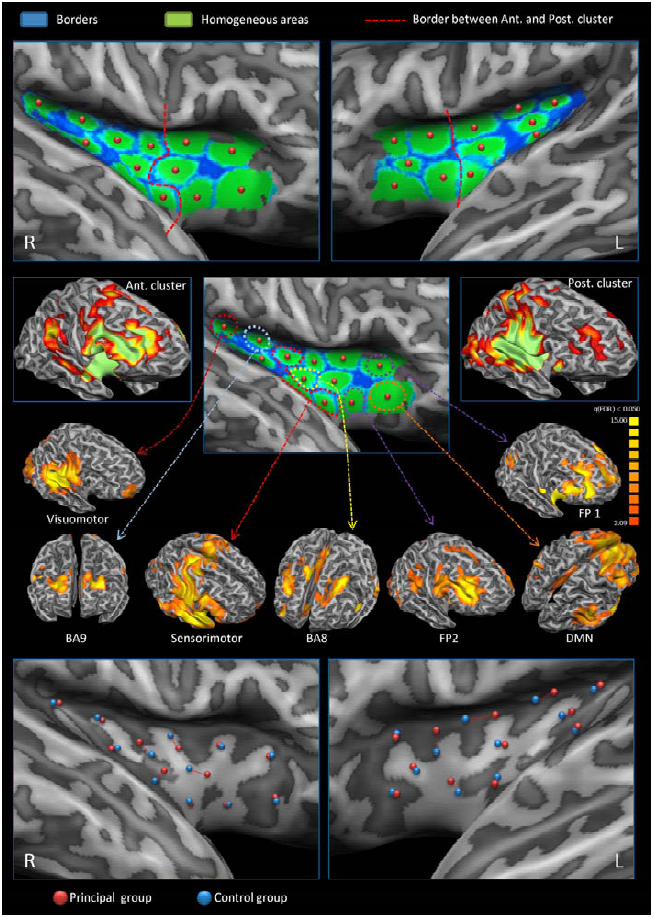
High-dimensional clusterization and node creation of the insular cortex. Upper panel: high-dimensional insular clusterization. Homogeneous areas are shown in green and borders in blue. The dotted red line outlines the separation between anterior and posterior clusters as detected in our previous studies (Cauda et al. 2011; Cauda et al. 2012b). Middle upper panels: anterior and posterior cluster functional connectivity. Middle lower panels: patterns of functional connectivity regressing out the common variance. Lower panels: Reliability of node creation. Nodes calculated from the main group are shown in red, nodes calculated from the replication group in blue. If separated, paired nodes are shown by a dotted red line.

## DISCUSSION

By applying a high-dimensional clustering procedure to the analysis of the functional connectivity of the insula, we were able to detect the connectivity patterns or, as defined by Leech et al. “echoes” (Leech et al. 2012) of other neural networks that constitute the hierarchical sub-parcellation of the anterior and posterior patterns of connectivity previously described: the ventral anterior cingulo-fronto-parietal “salience detection” network and dorsal posterior sensorimotor network, respectively. We successfully generated nodes, and the data obtained with this method were replicable with other datasets. As reported previously (Cauda et al. 2012a; Cauda et al. 2011; Cauda et al. 2012b; Kelly et al. 2012), there was some interhemispheric lateralization in the connectivity patterns of the insulae, indeed the localization of the clusters in the right and the left insulae was slightly different, especially in the posterior insula. These results were also confirmed when we subdivided the ROIs on the basis of their involvement in the two anterior and posterior clusters. A possible explanation for this hemispheric asymmetry involves some aspects of emotional and sympathetic processing, indeed the right insula responds more to sympathetic arousal whereas the left insula to parasympathetic nervous functions (Craig 2005; Harrison et al. 2010). Furthermore, the anatomical connectivity of the anterior insula with areas pertaining to the ventral attentional network, and in particular the temporo parietal junction (TPJ), has been demonstrated to be lateralized on the right side (Kucyi et al. 2012). This is coherent with the fight or flight function of the sympathetic system that requires an evaluation of the harmful potential of the incoming stimuli.

The posterior insula is characterized by smaller clusters and lesser homogeneity than the anterior. Consistently, hemispheric asymmetry is also more evident here than in the anterior insula. This finding supports the idea that the posterior circuits bear more heterogeneous connectivity patterns, in line with Craig’s hypothesis which suggests that the posterior insula can serve as a data collector from many different networks (Craig 2009, 2011). This group of posterior clusters have a more sensorimotor pattern of connectivity, as reported in some recent papers (Chang et al. 2012; Kelly et al. 2012). Indeed, moving from the more posterior part of the insular surface, the connectivity changes from visuomotor to prefrontal (BA9), then to sensorimotor and back to prefrontal (BA8) in the middle insular cortex. In fact not only action and perception-related but also some prefrontal patterns of connectivity are present in the posterior insula: this is explained by the involvement of this area in a variety of different activities other than action and perception such as pain, language, interoception, and sexuality as recently reported by some studies (Kelly et al. 2012; Kurth et al. 2010b). Overall, the posterior insula shows a more specific connectivity pattern than the anterior insula. The latter, conversely, shows connectivity with networks related to the switching of attention between internal and external-related stimuli such as attentional and default mode networks. Rather than being specific, this pattern suggests a function that tends to be a requirement of everyday activities. This has also been confirmed by several papers that have shown how the anterior insula is more aspecifically and massively activated in a broad series of different behavioral domains (Cauda et al. 2011; Chang et al. 2012; Kelly et al. 2012; Kurth et al. 2010b). In line with our data, these studies linked the activity of the anterior insula with cognitive and emotional responses: an involvement that, together with saliency detection (Seeley et al. 2007) and task switching (Dosenbach et al. 2007), is almost ubiquitous.

Some authors (Deen et al. 2011; Kurth et al. 2010b; Mutschler et al. 2009; Nelson et al. 2010b; Small 2010; Wager and Feldaman Barret 2004) have suggested a differentiation or a gradient of connectivity between the dorsal and ventral anterior insula that we failed to demonstrate in our previous papers. Indeed different levels of parcellation determined by the various methods used to calculate the optimal number of clusters lead to a different picture of the insular cortex. This phenomenon is particularly clear in the results of the study by Kelly et al. (Kelly et al. 2012) comparing insular parcellations with n=2 and n=3 clusters. In the parcellation with n=3 clusters the higher number of clusters made it possible to reveal an anterior ventral cluster that was not present with n=2. In the present paper the complex structure of the anterior insular cortex is further clarified: indeed we validated the recent identification by Touroutoglou et al. (Touroutoglou et al. 2012) of two dissociable frontoparietal patterns of functional connectivity in the anterior insular cortex, dorsal and ventral AI respectively, where a dorsal network (referred to here as dFP) and a ventral network (referred to here as vFP) coexist (for a similar result see also (Nelson et al. 2010b)). These two networks probably subserve two only partially different functions with the dorsal network involved more in the integration of top-down and bottom-up salient information and the ventral one in emotional salience detection aspects and the integration of bodily feelings (Touroutoglou et al. 2012). These two large-scale networks have a different pattern of connectivity: the dorsal network is centered more on dorsolateral and dorsomedial prefrontal cortices plus middorsal cingulate cortices, the ventral network is linked to the anterior cingulate, ventral prefrontal cortices and TPJ. Other authors have identified a network similar to the dorsal anterior cluster or to a mix of the dorsal and ventral anterior insular clusters, for example the frontoparietal control network (Vincent et al. 2008), the cingulo-opercular, ventral attentional (Corbetta et al. 2008; Shulman et al. 2009) and basal ganglia-frontoinsular (Shulman et al. 2009) control network (Dosenbach et al. 2007; Nelson et al. 2010b). Indeed the two anterior insular frontoparietal networks show areas of overlap and likely have a shared variance that in some conditions makes these two components less easily separable.

Interestingly, a cluster placed just in-between and anteriorly to these two areas shows connectivity with the Default Mode Network. Indeed this cluster is placed in a position that overlaps for a large part with the agranular area described by Mesulam and Mufson in the 1982 (Mesulam and Mufson 1982). This result seems to validate the hypothesis of an anterior insula arranged in a pivotal position and the continuous and variable reallocation of brain resources between internal and external focused networks (Spreng et al. 2010; Sridharan et al. 2008), modulating the switch between goal-oriented attentional and default mode networks. This fact would also explain the frequent activation of this brain area in such different tasks.

These small connectivity patterns that we demonstrated in our study and that further subdivide the anterior and posterior insular clusters are probably overcome by the variance of two other main patterns, but when the latter variances are regressed out, a more complex picture emerges. This behavior is explained by the hierarchical connectivity structure of the insular cortex as suggested by some recent papers (Cauda et al. 2012a; Kelly et al. 2012). Indeed, as also congruent with other studies (Kelly et al. 2012; Kurth et al. 2010a; Nanetti et al. 2009), the two clusters that were previously identified in our papers were here divided into a series of smaller parcels. This is also in line with the suggested intrinsically hierarchical structure of this area (Doucet et al. 2011; Power et al. 2011; Yeo et al. 2011) as hypothesized in the theories of Craig and Damasio (Craig 2005, 2009; Damasio 2005).

Therefore, the insula emerged as a much more complex structure than previously hypothesized, since echoes of various cerebral networks were revealed when we moved toward a more detailed parcellation of its surface.

In this study, we demonstrated that the detection of nodes using high-dimensional fuzzy c-mean parcellation is a simple, efficient and reliable method, which clearly demonstrates that the insula displays a hierarchical structure where information coming from the environment and from the body is integrated and distributed to different areas of the brain. Although our study confirmed the two (or three) major insular subdivisions, a more in-depth investigation showed that these areas can be further subdivided into smaller clusters each one characterized by its own pattern of connectivity that can be detected using appropriate techniques. Given the hierarchical structure of this area, moving the “cut line” through more detailed parcellations would reveal a more intricate architecture where it is possible to detect smaller areas with their own pattern of functional connectivity nested within bigger clusters. Indeed it can be hypothesized that, to exchange information with other areas (and thus with other networks), the different parts of the insula have to covariate with the “target areas”.

## References

Bandettini PA, Bullmore E (2008) Endogenous oscillations and networks in functional magnetic resonance imaging. Human brain mapping 29 (7):737–739. doi:10.1002/hbm.20607

Birn RM, Murphy K, Bandettini PA (2008) The effect of respiration variations on independent component analysis results of resting state functional connectivity. Human brain mapping 29 (7):740–750. doi:10.1002/hbm.20577

Bowman FD, Patel R, Lu C (2004) Methods for detecting functional classifications in neuroimaging data. Human brain mapping 23 (2):109–119. doi:10.1002/hbm.20050

Bullmore E, Sporns O (2009) Complex brain networks: graph theoretical analysis of structural and functional systems. Nature reviews Neuroscience 10 (3):186–198. doi:10.1038/nrn2575

Cauda F, Costa T, Torta DM, Sacco K, D’Agata F, Duca S, Geminiani G, Fox PT, Vercelli A (2012a) Meta-analytic clustering of the insular cortex: Characterizing the meta-analytic connectivity of the insula when involved in active tasks. NeuroImage. doi:10.1016/j.neuroimage.2012.04.012

Cauda F, D’Agata F, Sacco K, Duca S, Geminiani G, Vercelli A (2011) Functional connectivity of the insula in the resting brain. NeuroImage 55 (1):8–23. doi:10.1016/j.neuroimage.2010.11.049

Cauda F, Geminiani G, D’Agata F, Sacco K, Duca S, Bagshaw AP, Cavanna AE (2010) Functional connectivity of the posteromedial cortex. PloS one 5 (9). doi:10.1371/journal.pone.0013107

Cauda F, Torta DM, Sacco K, D’Agata F, Geda E, Duca S, Geminiani G, Vercelli A (2012b) Functional anatomy of cortical areas characterized by Von Economo neurons. Brain structure & function. doi:10.1007/s00429-012-0382-9

Cerliani L, Thomas RM, Jbabdi S, Siero JC, Nanetti L, Crippa A, Gazzola V, D’Arceuil H, Keysers C (2011) Probabilistic tractography recovers a rostrocaudal trajectory of connectivity variability in the human insular cortex. Human brain mapping. doi:10.1002/hbm.21338

Chang LJ, Yarkoni T, Khaw MW, Sanfey AG (2012) Decoding the Role of the Insula in Human Cognition: Functional Parcellation and Large-Scale Reverse Inference. Cereb Cortex. doi:10.1093/cercor/bhs065

Cloutman LL, Binney RJ, Drakesmith M, Parker GJ, Lambon Ralph MA (2012) The variation of function across the human insula mirrors its patterns of structural connectivity: evidence from in vivo probabilistic tractography. NeuroImage 59 (4):3514–3521. doi:10.1016/j.neuroimage.2011.11.016

Cohen AL, Fair DA, Dosenbach NU, Miezin FM, Dierker D, Van Essen DC, Schlaggar BL, Petersen SE (2008) Defining functional areas in individual human brains using resting functional connectivity MRI. NeuroImage 41 (1):45–57. doi:10.1016/j.neuroimage.2008.01.066

Corbetta M, Patel G, Shulman GL (2008) The reorienting system of the human brain: from environment to theory of mind. Neuron 58 (3):306–324. doi:10.1016/j.neuron.2008.04.017

Craig AD (2005) Forebrain emotional asymmetry: a neuroanatomical basis? Trends in cognitive sciences 9 (12):566–571. doi:10.1016/j.tics.2005.10.005

Craig AD (2009) How do you feel--now? The anterior insula and human awareness. Nature reviews Neuroscience 10 (1):59–70. doi:10.1038/nrn2555

Craig AD (2011) Significance of the insula for the evolution of human awareness of feelings from the body. Annals of the New York Academy of Sciences 1225:72–82. doi:10.1111/j.1749-6632.2011.05990.x

Damasio AR (2005) Descartes’ Error: Emotion, Reason, and the Human Brain.

Deen B, Pitskel NB, Pelphrey KA (2011) Three systems of insular functional connectivity identified with cluster analysis. Cereb Cortex 21 (7):1498–1506. doi:10.1093/cercor/bhq186

Dosenbach NU, Fair DA, Miezin FM, Cohen AL, Wenger KK, Dosenbach RA, Fox MD, Snyder AZ, Vincent JL, Raichle ME, Schlaggar BL, Petersen SE (2007) Distinct brain networks for adaptive and stable task control in humans. Proceedings of the National Academy of Sciences of the United States of America 104 (26):11073–11078. doi:10.1073/pnas.0704320104

Doucet G, Naveau M, Petit L, Delcroix N, Zago L, Crivello F, Jobard G, Tzourio-Mazoyer N, Mazoyer B, Mellet E, Joliot M (2011) Brain activity at rest: a multiscale hierarchical functional organization. Journal of neurophysiology 105 (6):2753–2763. doi:10.1152/jn.00895.2010

Fox MD, Zhang D, Snyder AZ, Raichle ME (2009) The Global Signal and Observed Anticorrelated Resting State Brain Networks. J Neurophysiol. doi:90777.2008[pii]10.1152/jn.90777.2008

Harrison NA, Gray MA, Gianaros PJ, Critchley HD (2010) The embodiment of emotional feelings in the brain. The Journal of neuroscience: the official journal of the Society for Neuroscience 30 (38):12878–12884. doi:10.1523/JNEUROSCI.1725-10.2010

Jakab A, Molnar PP, Bogner P, Beres M, Berenyi EL (2012) Connectivity-based parcellation reveals interhemispheric differences in the insula. Brain topography 25 (3):264–271. doi:10.1007/s10548-011-0205-y

Kelly C, Toro R, Di Martino A, Cox CL, Bellec P, Castellanos FX, Milham MP (2012) A convergent functional architecture of the insula emerges across imaging modalities. NeuroImage 61 (4):1129–1142. doi:10.1016/j.neuroimage.2012.03.021

Kiviniemi V, Starck T, Remes J, Long X, Nikkinen J, Haapea M, Veijola J, Moilanen I, Isohanni M, Zang YF, Tervonen O (2009) Functional segmentation of the brain cortex using high model order group PICA. Human brain mapping 30 (12):3865–3886. doi:10.1002/hbm.20813

Kucyi A, Moayedi M, Weissman-Fogel I, Hodaie M, Davis KD (2012) Hemispheric asymmetry in white matter connectivity of the temporoparietal junction with the insula and prefrontal cortex. PloS one 7 (4):e35589. doi:10.1371/journal.pone.0035589

Kurth F, Eickhoff SB, Schleicher A, Hoemke L, Zilles K, Amunts K (2010a) Cytoarchitecture and probabilistic maps of the human posterior insular cortex. Cereb Cortex 20 (6):1448–1461. doi:10.1093/cercor/bhp208

Kurth F, Zilles K, Fox PT, Laird AR, Eickhoff SB (2010b) A link between the systems: functional differentiation and integration within the human insula revealed by meta-analysis. Brain structure & function 214 (5-6):519–534. doi:10.1007/s00429-010-0255-z

Leech R, Braga R, Sharp DJ (2012) Echoes of the brain within the posterior cingulate cortex. The Journal of neuroscience: the official journal of the Society for Neuroscience 32 (1):215–222. doi:10.1523/JNEUROSCI.3689-11.2012

Leech R, Kamourieh S, Beckmann CF, Sharp DJ (2011) Fractionating the default mode network: distinct contributions of the ventral and dorsal posterior cingulate cortex to cognitive control. The Journal of neuroscience: the official journal of the Society for Neuroscience 31 (9):3217–3224. doi:10.1523/JNEUROSCI.5626-10.2011

Mesulam MM, Mufson EJ (1982) Insula of the old world monkey. I. Architectonics in the insulo-orbito-temporal component of the paralimbic brain. The Journal of comparative neurology 212 (1):1–22. doi:10.1002/cne.902120102

Miezin FM, Maccotta L, Ollinger JM, Petersen SE, Buckner RL (2000) Characterizing the hemodynamic response: effects of presentation rate, sampling procedure, and the possibility of ordering brain activity based on relative timing. NeuroImage 11 (6 Pt 1):735–759. doi:10.1006/nimg.2000.0568

Mutschler I, Wieckhorst B, Kowalevski S, Derix J, Wentlandt J, Schulze-Bonhage A, Ball T (2009) Functional organization of the human anterior insular cortex. Neuroscience letters 457 (2):66–70. doi:10.1016/j.neulet.2009.03.101

Nanetti L, Cerliani L, Gazzola V, Renken R, Keysers C (2009) Group analyses of connectivity-based cortical parcellation using repeated k-means clustering. NeuroImage 47 (4):1666–1677. doi:10.1016/j.neuroimage.2009.06.014

Napadow V, Dhond R, Conti G, Makris N, Brown EN, Barbieri R (2008) Brain correlates of autonomic modulation: combining heart rate variability with fMRI. NeuroImage 42 (1):169–177. doi:10.1016/j.neuroimage.2008.04.238

Nelson SM, Cohen AL, Power JD, Wig GS, Miezin FM, Wheeler ME, Velanova K, Donaldson DI, Phillips JS, Schlaggar BL, Petersen SE (2010a) A parcellation scheme for human left lateral parietal cortex. Neuron 67 (1):156–170. doi:10.1016/j.neuron.2010.05.025

Nelson SM, Dosenbach NU, Cohen AL, Wheeler ME, Schlaggar BL, Petersen SE (2010b) Role of the anterior insula in task-level control and focal attention. Brain structure & function 214 (5-6):669–680. doi:10.1007/s00429-010-0260-2

Power JD, Cohen AL, Nelson SM, Wig GS, Barnes KA, Church JA, Vogel AC, Laumann TO, Miezin FM, Schlaggar BL, Petersen SE (2011) Functional network organization of the human brain. Neuron 72 (4):665–678. doi:10.1016/j.neuron.2011.09.006

Rousseeuw PJ (1987) Silhouettes - a Graphical Aid to the Interpretation and Validation of Cluster-Analysis. J Comput Appl Math 20:53–65

Seeley WW, Menon V, Schatzberg AF, Keller J, Glover GH, Kenna H, Reiss AL, Greicius MD (2007) Dissociable intrinsic connectivity networks for salience processing and executive control. The Journal of neuroscience: the official journal of the Society for Neuroscience 27 (9):2349–2356. doi:10.1523/JNEUROSCI.5587-06.2007

Shulman GL, Astafiev SV, Franke D, Pope DL, Snyder AZ, McAvoy MP, Corbetta M (2009) Interaction of stimulus-driven reorienting and expectation in ventral and dorsal frontoparietal and basal ganglia-cortical networks. The Journal of neuroscience: the official journal of the Society for Neuroscience 29 (14):4392–4407. doi:10.1523/JNEUROSCI.5609-08.2009

Small DM (2010) Taste representation in the human insula. Brain structure & function 214 (5-6):551–561. doi:10.1007/s00429-010-0266-9

Smith SM, Miller KL, Moeller S, Xu J, Auerbach EJ, Woolrich MW, Beckmann CF, Jenkinson M, Andersson J, Glasser MF, Van Essen DC, Feinberg DA, Yacoub ES, Ugurbil K (2012) Temporally-independent functional modes of spontaneous brain activity. Proceedings of the National Academy of Sciences of the United States of America 109 (8):3131–3136. doi:10.1073/pnas.1121329109

Smith SM, Miller KL, Salimi-Khorshidi G, Webster M, Beckmann CF, Nichols TE, Ramsey JD, Woolrich MW (2011) Network modelling methods for FMRI. NeuroImage 54 (2):875–891. doi:10.1016/j.neuroimage.2010.08.063

Spreng RN, Stevens WD, Chamberlain JP, Gilmore AW, Schacter DL (2010) Default network activity, coupled with the frontoparietal control network, supports goal-directed cognition. NeuroImage 53 (1):303–317. doi:10.1016/j.neuroimage.2010.06.016

Sridharan D, Levitin DJ, Menon V (2008) A critical role for the right fronto-insular cortex in switching between central-executive and default-mode networks. Proceedings of the National Academy of Sciences of the United States of America 105 (34):12569–12574. doi:10.1073/pnas.0800005105

Talairach J, Tournoux P (1988) Co-planar stereotaxic atlas of the human brain: 3-dimensional proportional system: an approach to cerebral imaging. Thieme,

Taylor KS, Seminowicz DA, Davis KD (2009) Two systems of resting state connectivity between the insula and cingulate cortex. Human brain mapping 30 (9):2731–2745. doi:10.1002/hbm.20705

Touroutoglou A, Hollenbeck M, Dickerson BC, Feldman Barrett L (2012) Dissociable large-scale networks anchored in the right anterior insula subserve affective experience and attention. NeuroImage 60 (4):1947–1958. doi:10.1016/j.neuroimage.2012.02.012

Vincent JL, Kahn I, Snyder AZ, Raichle ME, Buckner RL (2008) Evidence for a frontoparietal control system revealed by intrinsic functional connectivity. Journal of neurophysiology 100 (6):3328–3342. doi:10.1152/jn.90355.2008

Wager TD, Feldaman Barret L (2004) From affect to control: Functional specialization of the insula in motivation and regulation. online PsycExtra: http://wwwcolumbiaedu/cu/psychology/tor/

Yeo BT, Krienen FM, Sepulcre J, Sabuncu MR, Lashkari D, Hollinshead M, Roffman JL, Smoller JW, Zollei L, Polimeni JR, Fischl B, Liu H, Buckner RL (2011) The organization of the human cerebral cortex estimated by intrinsic functional connectivity. Journal of neurophysiology 106 (3):1125–1165. doi:10.1152/jn.00338.2011

Zuo XN, Kelly C, Adelstein JS, Klein DF, Castellanos FX, Milham MP (2010) Reliable intrinsic connectivity networks: test-retest evaluation using ICA and dual regression approach. NeuroImage 49 (3):2163–2177. doi:10.1016/j.neuroimage.2009.10.080

